# Prediction of risk scores for colorectal cancer patients from the concentration of proteins involved in mitochondrial apoptotic pathway

**DOI:** 10.1101/639740

**Authors:** Anjali Lathwal, Chakit Arora, Gajendra P. S. Raghava

## Abstract

One of the major challenges in managing the treatment of colorectal cancer (CRC) patients is to predict risk scores or level of risk for CRC patients. In past, several biomarkers, based on concentration of proteins involved in type-2/intrinsic/mitochondrial apoptotic pathway, have been identified for prognosis of colorectal cancer patients. Recently, a prognostic tool DR MOMP has been developed that can discriminate high and low risk CRC patients with reasonably high accuracy (Hazard Ratio, HR = 5.24 and p-value = 0.0031). This prognostic tool showed an accuracy of 59.7% when used to predict favorable/unfavorable survival outcomes. In this study, we developed knowledge based models for predicting risk scores of CRC patients. Models were trained and evaluated on 134 stage III CRC patients. Firstly, we developed multiple linear regression based models using different techniques and achieved a maximum HR value of 6.34 with p-value = 0.0032 for a model developed using LassoLars technique. Secondly, models were developed using a parameter optimization technique and achieved a maximum HR value of 38.13 with p-value 0.0006. We also predicted favorable/unfavorable survival outcomes and achieved maximum prediction accuracy value of 71.64%. The performance of our models were evaluated using five-fold cross-validation technique. For providing service to the community we also developed a web server ‘CRCRpred’, to predict risk scores of CRC patients, which is freely available at https://webs.iiitd.edu.in/raghava/crcrpred.

## Introduction

Colorectal cancer (CRC) or large bowel cancer is one of the prevalent and fatal cancers with about 95 percent of them as adeno-carcinomas [1]. It is the third most widely diagnosed cancer in males and the second in females, with 1.84 million new cases and roughly about 883,200 deaths in 2018 [1]. A comprehensive information about CRC is available in WHO’s database GLOBOCAN that includes worldwide incidence, country-specific incidence, and death rates [1]. In the United States alone, around 146,600 new instances of colorectal cancer are estimated for the year 2019, of which 101,420 will be colon, and the rest will be rectal tumors [2]. Approximately 51,020 Americans are expected to succumb due to CRC in 2019 [2], representing roughly 8 percent of all cancer deaths. The frequency of CRC, internationally ranges between 6-8 folds with the most prominent incidence rates in Australia, New Zealand, Europe, and North America, and the least rates in Africa and South-Central Asia [1]. These geographic contrasts seem, by all accounts, to be inferable from the disparity in dietary and ecological exposures that are forced upon a background of genetically determined susceptibility. While, environmental factors (such as lifestyle, diet, physical activity, etc.) have been reported to play an important role [3, 4], several genetic/epigenetic changes have been elucidated to cause/promote CRC in the past decades [5]. A significant number of biological experiments along with several theoretical/bioinformatics studies based on genomics/proteomics data have revealed a crucial role of the mitochondrial apoptotic pathway in tumor progression [6, 7].

Mitochondrial or apoptotic type 2 signaling pathway is a conserved pathway in many organisms, which controls the lifespans of cells in different tissues and leads to cellular death or apoptosis as a result of genotoxic and other stresses [8]. Alteration in the expression of signaling proteins involved in this pathway has been associated with tumor survival/progression and chemo-resistance development [6, 7]. Apoptotic type 2 (intrinsic) pathway consists of a cascade of Bcl2 family proteins which are broadly classified into two categories: Anti-apoptotic proteins that include Bcl2, BclXL, Mcl1 and pro-apoptotic proteins that include Bax, Bak, Bid, Bim [9]. Each of these has a definite functional role in regulating the process of mitochondrial pore formation (Mitochondrial Outer Membrane Permeabilization) leading to activation of caspases and ultimately, the demise of the cell [8] (Fig 1). While anti-apoptotic proteins are agonists to effector pro-apoptotic proteins (such as Bak/Bax), BH3 only activator pro-apoptotic proteins such as Bid, Bim etc. can either cleave Bax/Bak to their active counterparts (exposed TM domains) which further oligomerize on mitochondrial membrane to form pores or, can bind to anti-apoptotic proteins and inhibit their function. Both of these scenarios lead to increased apoptotic activation. Dysregulation in the concentration of these proteins (mainly upregulation of anti-apoptotic proteins) benefits tumor cells in their survival and cancer development [10].

**Fig 1.**
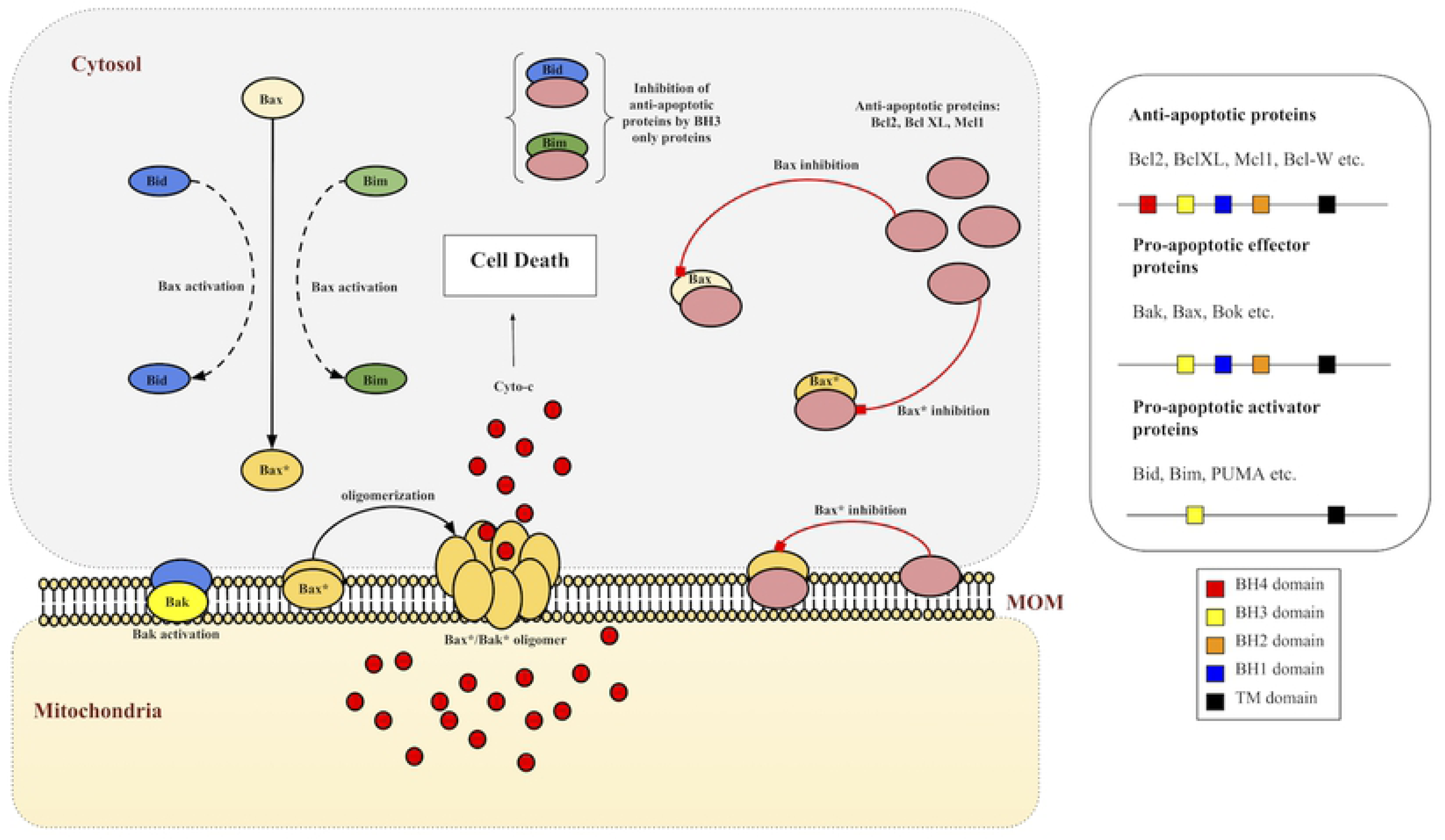
Mitochondrial type 2 pathway and MOMP. Bax/Bak proteins are activated by the action of BH3 only proteins Bid/Bim. Activated Bax/Bak oligomerize to form pores on mitochondrial outer membrane (MOM) as a result of which apoptotic factors such as cyto-c are released, a process known as mitochondrial outer membrane permeabilization (MOMP), and lead the cell towards death. Anti-apoptotic proteins such as Bcl2, BclXL and Mcl1 bind with both BH3 only proteins as well as pro-apoptotic Bax/Bak to inhibit their function.

This biological understanding has resulted in the design and development of several therapeutic strategies (and drugs) which target Bcl2 family proteins and exploit the mitochondrial pathway to induce apoptosis in tumor cells. Although, it has been observed that the failure rate of these chemotherapeutic drugs (due to tumor relapses) is significant, possibly due to variation in protein expressions [9] and/or variation in ligand-receptor binding affinities amongst proteins (due to mutations) within CRC patients [11, 12]. In 2013, a systems based model DR MOMP was introduced which incorporated concentration data of pro- and anti-apoptotic proteins to predict minimal dose response (BH3 only stress) required for MOMP, which was denoted as *η*. The authors in this study demonstrated that *η* can be used to classify a group of 26 CRC patients into responders (favorable outcomes) and non-responders (unfavorable outcomes) to chemotherapy [13]. Recently, application of model DR MOMP has been shown in classification of 134 chemotherapy-treated stage III CRC patients into high and low risk groups. It was observed that high risk patients classified by DR MOMP had around five-fold increased risk of death (HR=5.2, p-value=0.02) as compared to low risk patients [14]. It was successful in differentiation of high and low risk patients with an overall accuracy of 59.7%.

In this study, we made a systematic attempt to develop models for predicting risk score for CRC patients, which can be used to discriminate high and low risk patients. In contrast to DR MOMP, our models are data-driven where parameters are optimised from the protein concentration obtained from CRC patients. In order to evaluate performance of our models, we compute performance in terms of standard parameters such as Hazard ratio (HR) and Confidence Interval (CI). It is important to compare performance of models developed in this study using previously developed models such as DR MOMP. One of the objective of this study is to provide service to society, thus a web server, ‘CRCRpred’ (https://webs.iiitd.edu.in/raghava/crcrpred) has been developed that can predict risk score for CRC patients.

## Materials and methods

### Dataset

The dataset ‘CRC stage III cohort’ used in this study was obtained from [14]. It contains data from Formalin-fixed paraffin-embedded (FFPE) primary tumour samples from 134 patients treated with FOLFOX and XELOX chemotherapy regimens. Primarily, it includes Bcl2 family protein expression in nM extracted by reverse phase protein array. In addition, the dataset also contains complete clinical information such as survival time, censoring data, metastatic staging; obtained from medical monitoring of the patients.

### Survival Analysis

Computation of Hazard Ratios and CIs were carried out to predict the risks of death associated with high-risk and low-risk groups stratified on the basis of mean and median values of various histopathological factors, using the univariate unadjusted Cox proportional hazard models. Multivariate Cox proportional models were used to analyse multiple covariates further, to assess the relative death risks associated with different factors. Kaplan-Meier plots were used to compare survival curves of high risk and low risk groups [15]. Survival analyses on these datasets were performed using ‘survival’ package (V.2.42-6) in R (V.3.4.4, The R Foundation). Statistical significance between the survival curves were estimated using log-rank tests. Wald tests were performed to estimate significance of the explanatory variables used for HR calculations.

### Five-fold cross-validation

The dataset is shuffled randomly and divided into 5 groups and an iterative process begins. During each iteration, a unique group is taken as a test dataset and combination of remaining groups as a train dataset. Model is fitted on the train dataset and evaluated on the test dataset. Model’s performance are evaluated using standard parameters. The process is repeated 5 times and each sample is processed once as a testing data point and 4 times as training data point.

### Multiple linear regression

Multiple linear regression models from Python’s scikit-learn (v0.20.3) were implemented to fit the protein concentrations (independent variables) against the overall survival time (target variable), which can be represented as *OS* = *x*_0_ + *x*_1_*Bak* + *x*_2_*Bax* + *x*_3_*Bcl*2 + *x*_4_*BclXL* + *x*_5_*Mcl*1. Simple linear regression (ordinary least squares), ridge regression, lasso regression, lasso lars regression, bayesian ridge regression and elastic net regression methods are the techniques that were used to estimate the coefficients *x*_0_, *x*_1_, .. *x*_5_. The fitting and test evaluations were carried using a five-fold cross-validation scheme. Combination of all five evaluated test datasets (predicted OS) was then used to classify the actual patient survival time (OS) at mean and median cutoffs to estimate HR, CI and p-values. Coefficient optimization and regularization was achieved using the in-built methods such as RidgeCV, LassoCV, LassoLarsCV etc.

### Parameter optimization technique and estimation of Risk Score (RS)

A column vector *β* was defined as *β* = *XW ^T^* where, *W* = (*w*_*Bak*_ *w*_*Bax*_ *w*_*Bcl*2_ *w*_*BclXL*_ *w*_*Mcl*1_), is a 1 *×* 5 weight vector containing optimized coefficients *w*_*k*_ ∈ [-1, 1] and X is a *n ×* 5 dataset matrix where n rows signify patients in the dataset and 5 columns have the protein expression values s.t for a patient j and at *W* = *W*_*i*_:

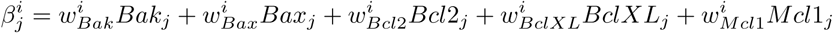

The dataset corresponding to 134 stage III CRC patients was uniformly divided into 5 subsets. A parallel parameter (W) search upto one decimal digit precision was performed iteratively, in order to maximize the objective ‘Hazard Ratio’ at mean and median cutoffs, on five training sets (each train set ∼ 4 out of 5 subsets). Running five search algorithms parallely with reduced yet homogeneous data saved computation time. The top parameter set obtained for each case (W1,W2,…W5) was then used to evaluate *β*^*i*^ and estimate HR values on the complete dataset using mean and median cutoffs. Algorithm is given in the Fig 2. For precision upto two decimal digits, W* was evaluated as:

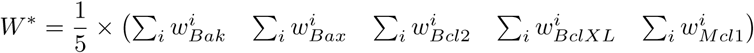

where i = 1,2,…5. From this point onwards we will refer the z-normalized version of the vector *β*^*\**^ as Risk Score (RS), for reasons that will become clear later.

**Fig 2.**
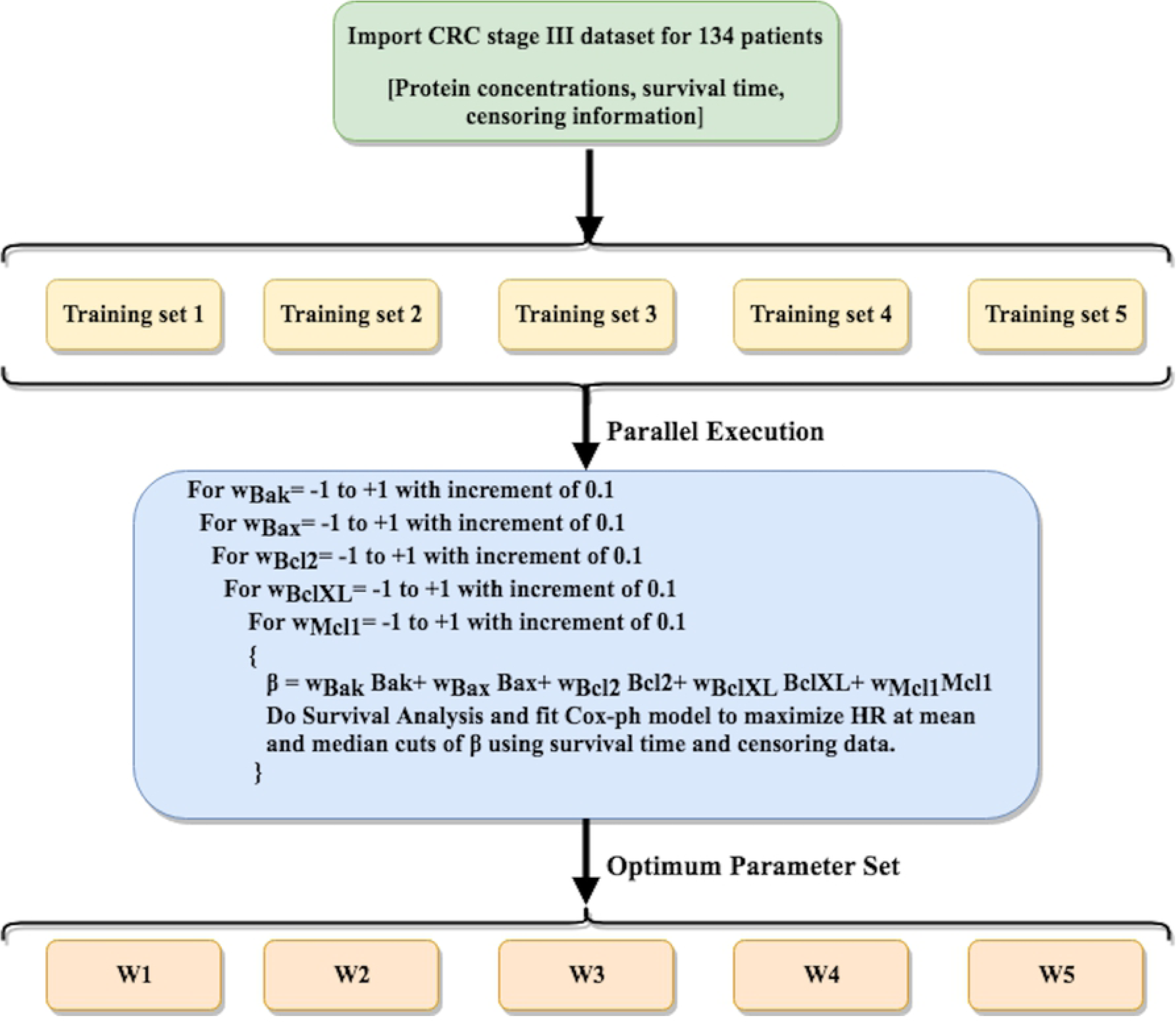
Algorithm for the parameter optimization technique. Pseudocode for parallel parameter optimization on five training sets in order to maximize HR as objective function.

## Results

### Single variable based classification

Following the univariate analysis presented in [14], we stratified high and low risk patients on the basis of various factors at median cutoff. The results in Table 1 show the HR, CI and p values due to these factors. We estimated hazard ratios and CIs on the basis of each protein concentration to examine whether any of them can act as a clinical marker that differentiates high risk and low risk CRC patients. As shown in Table 1, HR varies from 1.3 (Age) to 20.87 (BclXL). Out of these, BclXL was able to differentiate high and low risk CRC patients, on the basis of mean (HR=7.19, p-value=0.0004) [14] and median (HR=20.81, p=0.0030) cutoffs, thus achieving maximum differentiation. To establish BclXL as an exclusive prognostic marker, we calculated differences in mean levels of all the proteins between patients that survived the study and patients who succumbed to death or whose cancer relapsed. A t-test was performed for each of these proteins, and it was observed that levels of Bak (p=0.0042), Bax (p=0.0094), BclXL (p=3.5e-05) and Mcl1 (p=0.02) were significantly different between the two groups. This result indicated the importance of other proteins in combination with BclXL and rejected the possibility of using BclXL as the only biomarker. As a result, total protein levels (Bcl2+BclXL+Mcl1+Bax+Bak) and difference between anti- and pro-apoptotic protein levels (Bcl2+BclXL+Mcl1-Bax-Bak), were also used to estimate HR values. These showed good results in the case of CRC dataset. Total protein concentration was able to differentiate high and low risk patients with HR=5.81, p-value=0.0010 at mean cutoff (not shown here) and HR=6.37, p-value=0.0030 at median cutoff. While difference between anti- and pro-apoptotic protein levels was able to differentiate high and low risk patients with HR=3.05, p-value=0.02 at mean cutoff (not shown here), it was insignificant at median cutoff (p-value*<* 0.05).

**Table 1.** Univariate analysis for CRC stage III dataset. Survival analysis on the basis of all possible numeric features was performed to estimate HR and CIs on the basis of stratification of patients via median cut.

| CRC stage III cohort (n=134) | > Median(Factor) |  |
| --- | --- | --- |
| Factor | HR(95% CI) | p-value |
| 1. Age | 1.3 (0.54 - 3.14) | 0.5500 |
| 2. Bax | 1.34 (0.55 - 3.23) | 0.5200 |
| 3. Bak | 2.79 (1.07 - 7.3) | 0.0400* |
| 4. Bcl2 | 1.25 (0.51 - 3.02) | 0.6200 |
| 5. BclXL | 20.81 (2.7 - 155.5) | 0.0030* |
| 6. Mcl1 | 1.64 (0.67 - 4.03) | 0.2700 |
| 7. Bcl2+BclXL+Mcl1+Bax+Bak | 6.37 (1.86 - 21.73) | 0.0030* |
| 8. Bcl2+BclXL+Mcl1 - Bax - Bak | 2.49 (0.95 - 06.47) | 0.0600 |

### Multiple Linear Regression models for risk estimation

In order to find a relationship between level of protein concentration and survival time, we developed multiple linear regression models. The protein concentrations were used as independent/input variables and overall survival time (OS) was used as output/target/dependent variable. This implementation was done using Python’s sklearn package. After model fits and test validations, predicted OS from different methods was used to stratify high and low risk patients. Results are provided in Table 2.

**Table 2.** Multiple linear regression models for risk estimation. Evaluation of hazard ratios on CRC sample using various regression models with protein concentrations as features and overall survival time (OS) as target variable. LassoLars was found to be the best regression model for which HR was found to be maximum (HR=6.34,p=0.0032) for patients with predicted OS*<* median (predicted OS) estimated at higher risk of mortality.

| Model | < *Mean(OS)* |  | < *Median(OS)* |  |
| --- | --- | --- | --- | --- |
| Name | HR | p-value | HR | p-value |
| LR | 3.19 | 0.0132* | 3.27 | 0.0219* |
| Ridge | 3.34 | 0.0101* | 3.27 | 0.0219* |
| Lasso | 1.79 | 0.196 | 2.09 | 0.1170 |
| LassoLars | 2.44 | 0.0472* | 6.34 | 0.0032* |
| Elastic net | 1.79 | 0.1960 | 2.15 | 0.1030 |
| Bayesian ridge | 2.08 | 0.1010 | 2.64 | 0.0469* |

It was observed that LassoLars (LassoLarsCV from sklearn.linear model) based model performed better than other models and achieved a maximum HR value of 6.34 with p-value=0.0032 for predicted OS at median cutoff. KM plot for this is shown in Fig 3. Even though this method uses information about other proteins and provides predicted OS as a prognostic marker which performs better that many previously established markers, yet BclXL alone continues to be a better prognostic biomarker in the context of HR value.

**Fig 3.**
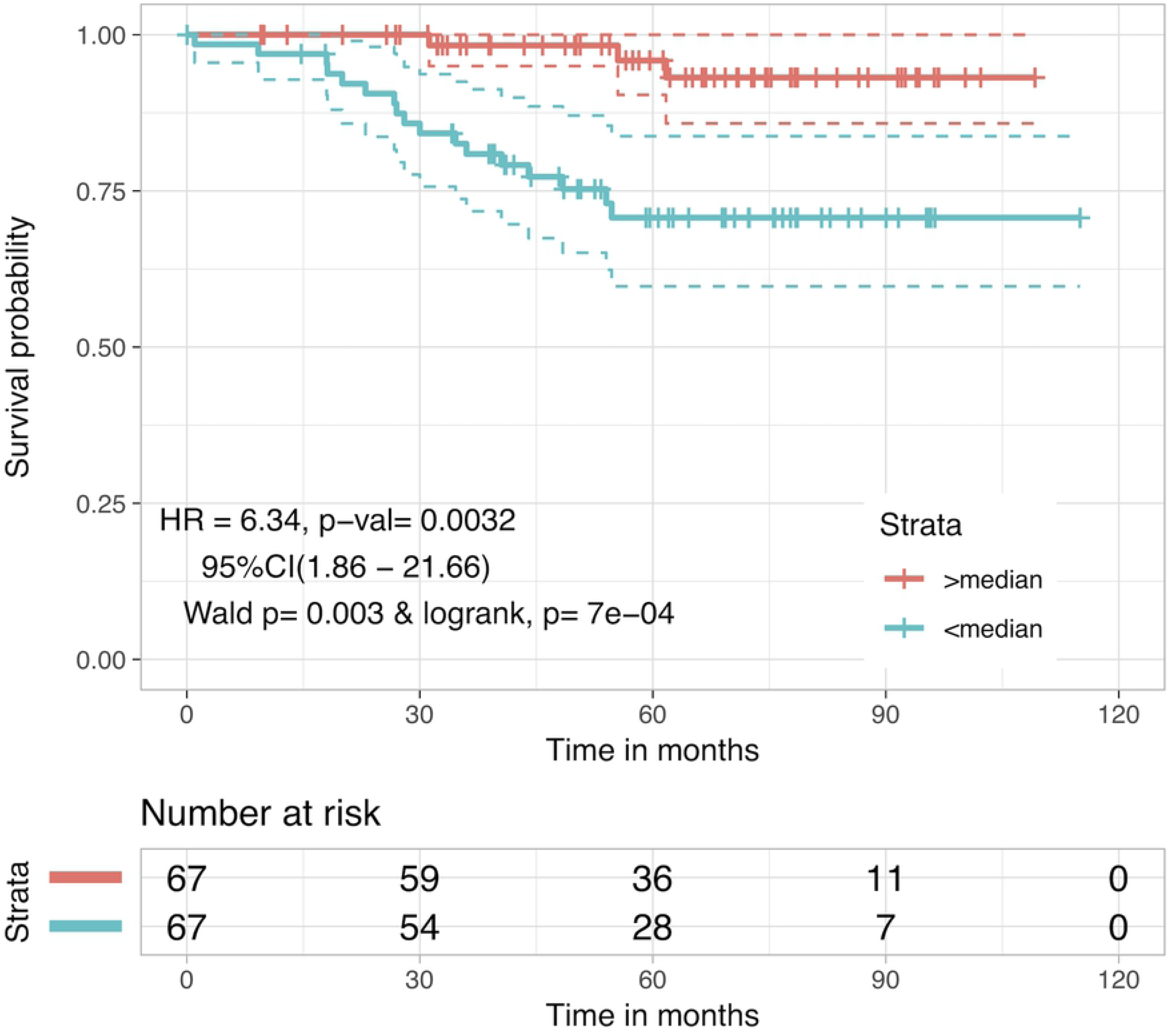
Kaplan Meier plot for risk stratification using LassoLars predicted OS. Patients were stratified using predicted OS estimated from LassoLars regression method after five-fold cross-validation. Patients with predicted OS*<*median(predicted OS) were at 6-fold higher risk than patients with OS*>*median(predicted OS) (HR=6.34, p-value=0.0032).

### Prediction of risk score using parameter optimization technique

One of the limitations of multiple linear regression models is that they are extensions of linear regression models where they evaluate relationship between a dependent and each independent variable. In addition, it is assumed that these protein concentrations (independent variables) have no relationship with each other. Thus, there is a need to develop a model which can handle non-linear data and correlated variables. In this study, we used a simple parameter optimization technique, where all possible weights were tried in an iterative manner to obtain best weights. By means of this method, Risk Score (RS) is predicted that is derived from anti and pro-apoptotic protein levels.

In parameter optimization technique, parameters or weights are optimized using iterative techniques which increases the possibility of over optimization. In order to avoid any over optimization, we used the concept of five-fold cross validation (Fig 2). Results corresponding to different parameter sets W1,W2…W5 and W* are given in Table 3. Patients with RS *<* 0 (mean) and RS *<* 0.266 (median), are found to be at higher risk with HR=38.13 (p-value=0.0004) and 22.27 (p-value=0.0025) respectively, than patients with RS *≥* 0 and RS *≥* 0.266. Kaplan Meier plots for this case are shown in Fig 4 (for other cases see Supplementary S1 File). It is evident from these results that while Bak, Bcl2 and Bax are somewhat less relevant for prognostic studies, BclXL and Mcl1 on the other hand, are the two dominating proteins to look at while stratifying CRC patients, due to their high weights (contribution) in *β*^*\**^. These results also correlate with isolated studies on BclXL and Mcl1 which showed their relevance as prognostic markers in the past [16, 17]. However, a biomarker such as RS accounts for a more comprehensive apoptotic profile and it makes more sense to use it as compared to any single protein.

**Table 3.** Evaluation of hazard ratios for each of the five parameter sets obtained from the train cases. *W*^*\**^ is the average of *W*_1_ - *W*_5_.

| Case | $< Mean(\beta)$ | | $< Median(\beta)$ | |
| --- | --- | --- | --- | --- |
| $W_i = (w_{Bak}^i \quad w_{Bax}^i \quad w_{Bcl2}^i \quad w_{BclXL}^i \quad w_{Mcl1}^i)$ | HR | p-value | HR | p-value |
| $W_1 = (0.0 \quad 0.2 \quad -0.1 \quad -0.8 \quad -0.9)$ | 33.23 | 0.0006 | 22.96 | 0.0023 |
| $W_2 = (0.0 \quad 0.1 \quad -0.1 \quad -0.9 \quad -0.3)$ | 18.88 | 8e-05 | 22.96 | 0.0023 |
| $W_3 = (0.0 \quad 0.2 \quad 0.0 \quad -0.9 \quad 0.0)$ | 15.94 | 0.0002 | 21.54 | 0.0028 |
| $W_4 = (0.0 \quad 0.2 \quad -0.2 \quad -0.9 \quad -0.8)$ | 11.26 | 0.0001 | 22.41 | 0.0024 |
| $W_5 = (0.1 \quad 0 \quad -0.1 \quad -0.7 \quad -0.7)$ | 11.03 | 0.0001 | 10.35 | 0.0017 |
| $W^* = (0.02 \quad 0.14 \quad -0.1 \quad -0.84 \quad -0.54)$ | 38.13 | 0.0004 | 22.27 | 0.0025 |

**Fig 4.**
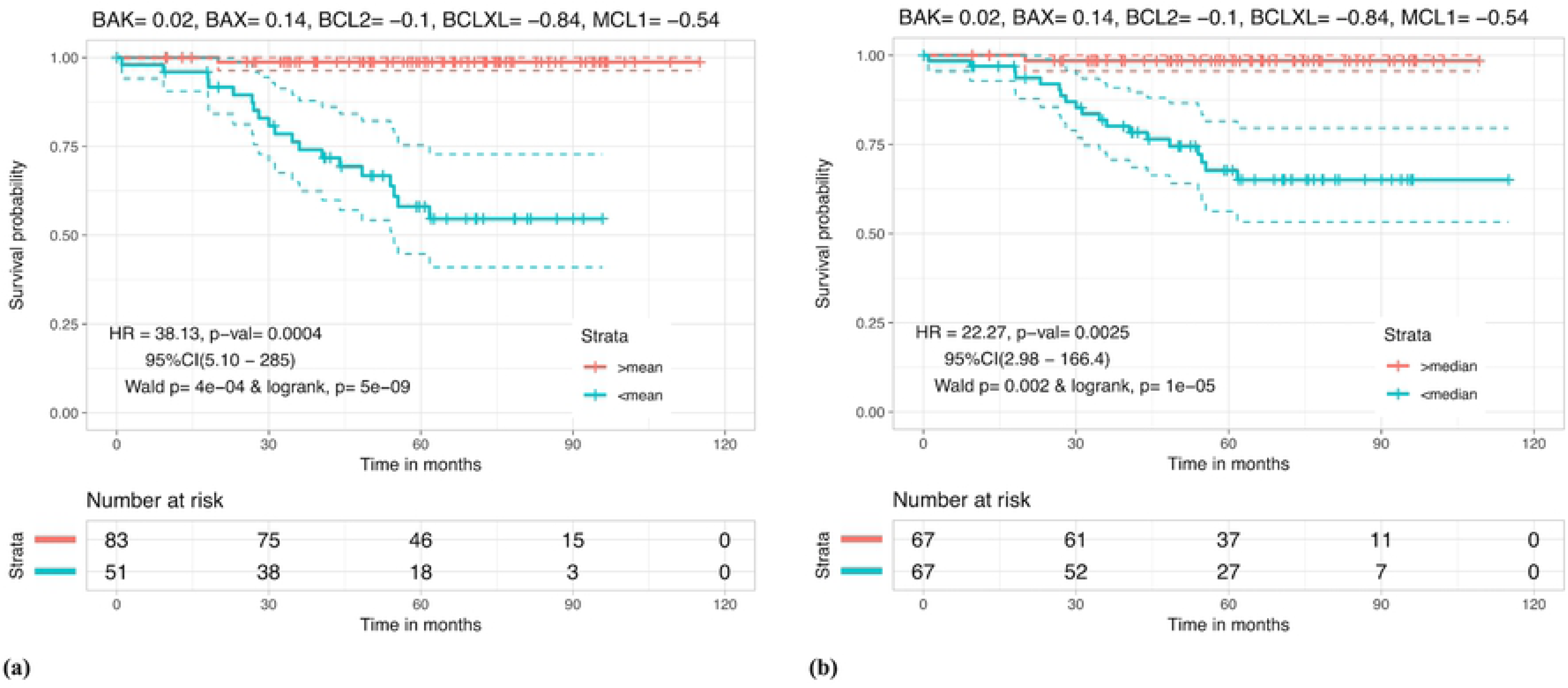
Kaplan Meier plot for risk stratification using Risk Score (RS). Kaplan Meier survival curves for risk estimation in the CRC patients based on the mean (RS=0) and median (RS=0.266) cutoffs. (a) Patients with RS *<* 0 were at around 38 fold greater risk relative to patients with RS *≥* 0 (HR = 38.13 and p-value = 0.0004). (b) Patients with RS *<* 0.266 were at around 22 fold greater risk relative to patients with RS *≥* 0.266 (HR = 22.27 and p = 0.0025).

### Prediction of favourable and unfavourable outcomes: a comparative study

The CRC stage III cohort dataset contains 95 favorable patients i.e patients that were alive during the 5 years study period and 39 unfavourable patients consisting of cases of recurrences/deaths. A comparison between a recently established mathematical model based prognostic marker, _*z*_*η* [14], and RS was performed on the basis of prediction accuracy of favorable/unfavorable outcomes when concentrations of apoptotic family proteins are known. RS showed a prediction accuracy of 71.64% at mean cutoff, as compared to 59.7% of _*z*_*η*. Results are summarized in Fig 5.

**Fig 5.**
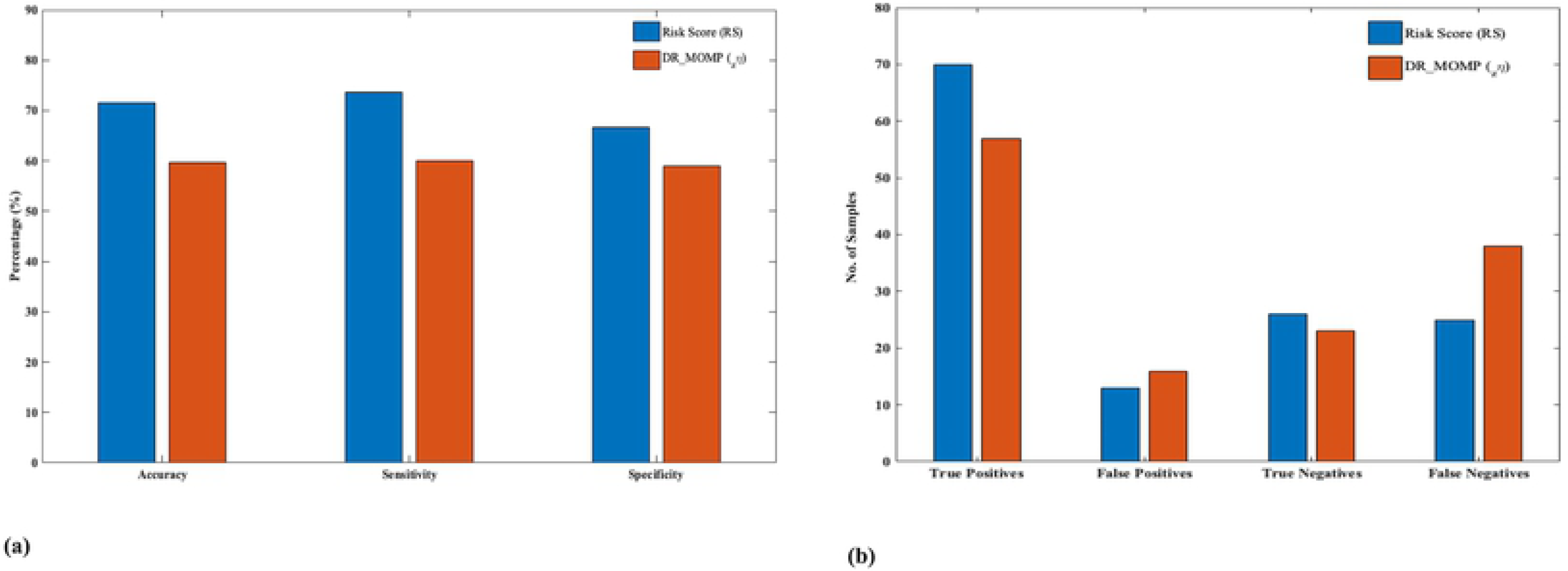
Kaplan Meier plot for risk stratification using Risk Score (RS). Prediction of favorable/unfavorable outcomes: (a) RS, at mean cutoff, is shown to predict favorable and unfavorable outcomes with an increased sensitivity (73.68%), specificity (66.66%) and accuracy (71.64%) as compared to DR MOMP’s _*z*_*η* (sens=60%, spec=58.9%, acc=59.7%). (b) A corresponding improvement in predicted positive (favorable) and negative (unfavorable) outcomes and reduced false predictions, by implementing RS, is presented here. (Favorable: RS*≥*0, Unfavorable: RS*<*0).

### Multivariate analysis reveals RS as the most significant factor associated with patient survival times

A multivariate analysis using cox proportional hazard models, was performed to see the association of other pathological features present in the dataset with the mortality risk of patients. RS was estimated for CRC stage III cohort dataset, to stratify high risk (RS *<* 0 or *<* 0.266) and low risk (RS *≥* 0 or *≥* 0.266) patients. Results are reported in Fig 6, clearly indicating that RS outperforms any other variable, when it comes to differentiation of patients on the basis of OS. RS is found to be associated with around 30 fold increased death risk in high risk patients as compared to low risk patients in CRC dataset (HR= 29.44, p-value= 0.001) for mean cutoff. It was also observed that *β*^1^, *β*^2^, *β*^3^, *β*^4^, *β*^5^ were also significant in a multivariate setting (results in Supplementary S1 File) as compared to other clinical factors, however RS (or *β*^*\**^) performs better than them. These results strengthen the conclusion that RS is a significantly improved prognostic marker over previously established histo-pathological biomarkers.

**Fig 6.**
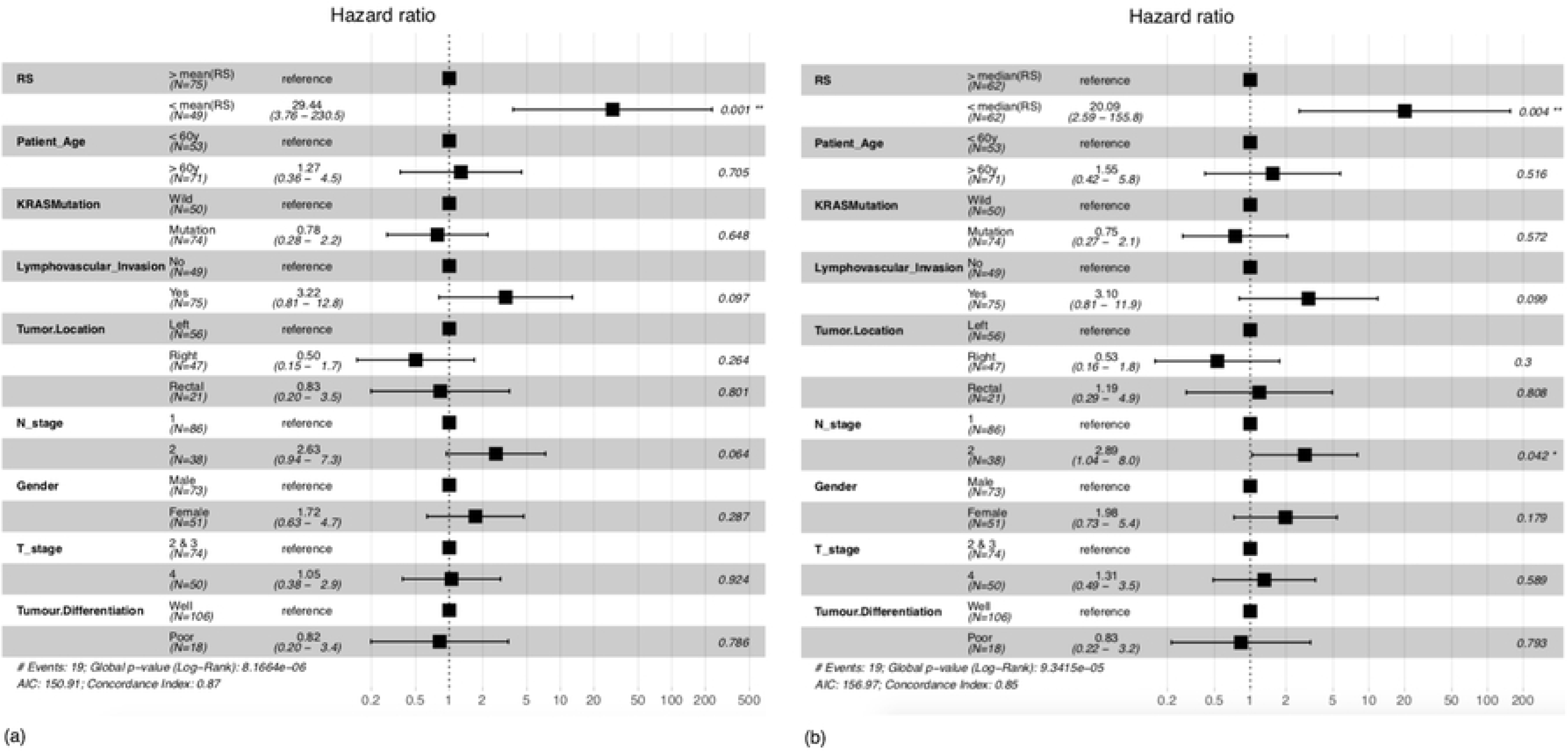
Multivariate COX-PH hazard analysis for risk estimation in the CRC patients based on RS. (a) At mean cutoff (RS=0). and (b) At median (RS=0.266) cutoff. Here right tumor location - hepatic flexure, caecum, ascending, traverse; left tumor location - splenic flexure, descending, sigmoid and rectal - rectum, rectosigmoid.

### RS differentiates high and low-risk patients belonging to subgroups formed on the basis of clinical/pathological features

Other factors like age [18], gender [18], TNM staging [19], lymphovascular invasion [20] and location of tumor in colon [21] have been shown to affect colorectal cancer incidence with preferable bias towards certain groups for e.g. incidence rates have been shown to be greater in males than females [18]. Based on these and the results from multivariate analysis, we looked at different sub-populations with a certain clinical/pathological feature in common. Using RS at mean and median cutoffs, we were able to stratify these sub-populations in CRC dataset into high risk and low risk groups. Results in the form of Kaplan Meier plots and logrank tests in Fig 7, show significant stratification of literature-established high risk sub-groups such as patients with older age [18], patients with tumor located in the right side (right tumor location [21]), patients with positive lymphovascular invasion [20], male patients [18], patients with tumor spread into lymph nodes [19] and patients with larger tumor sizes [19] into further high and low risk patients. Results for other subgroups and median cutoffs are given in Supplementary S1 File.

**Fig 7.**
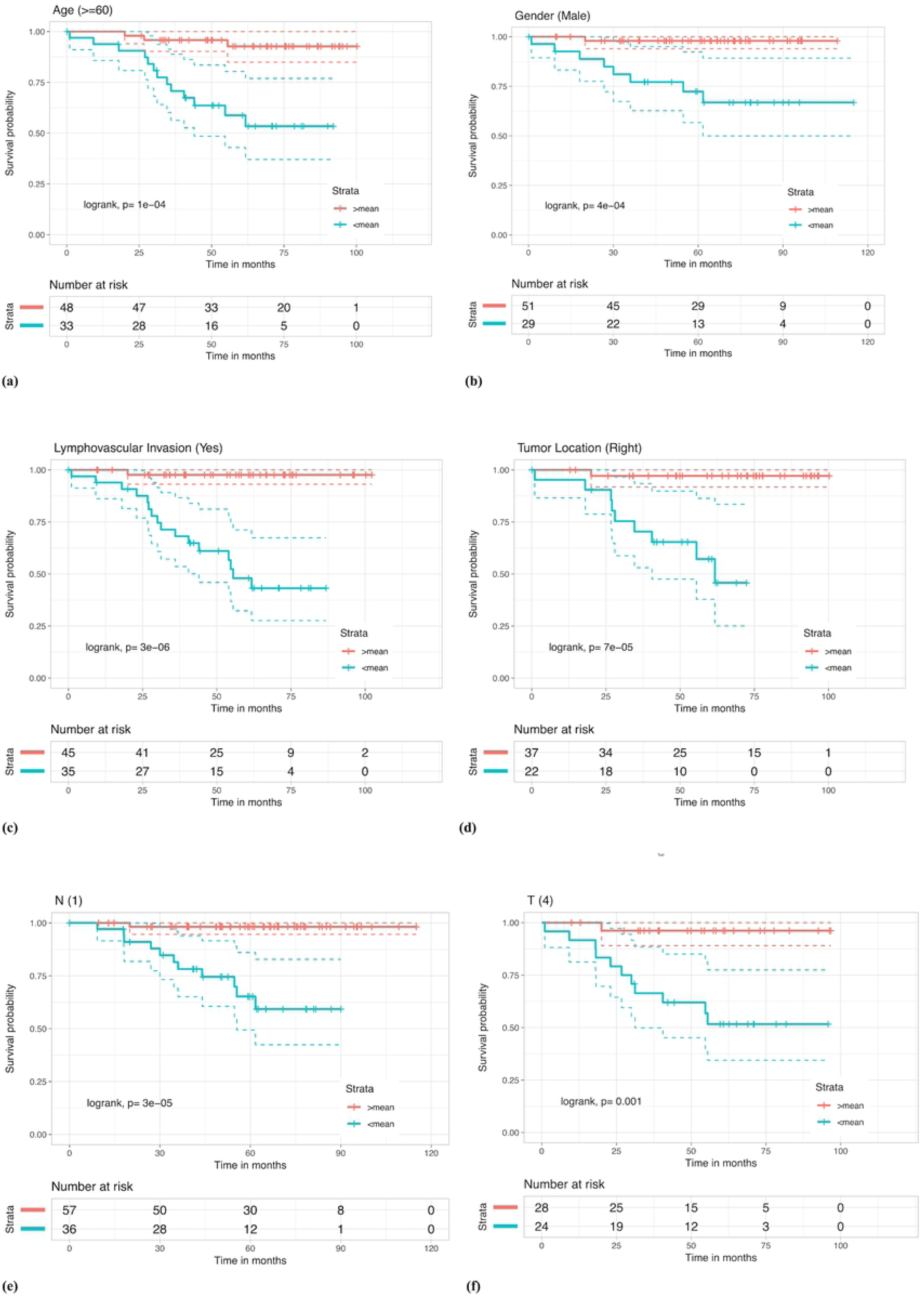
Kaplan Meier survival curves for risk estimation in the sub-populations of CRC stage III patient cohort based on RS using the mean (RS=0) cutoff, show significant differences in high/low risk groups. (a) Samples with patient age greater than 60y were stratified (log rank, p= 1e-04) in high/low risk groups with HR= 8.03 and p-value= 0.0017. (b) Male patients were stratified (log rank, p= 4e-04) within high/low risk groups with HR= 15.91 and p-value= 0.0091. (c) Patient with lymphovascular invasion were stratified (log rank, p= 3e-06) in high/low risk groups with HR= 24.92 and p-value= 0.0018. (d) Patients with tumor location in the right side were stratified (log rank, p= 7e-05) in high/low risk groups with HR= 20.16 and p-value= 0.0046. (e) Patient samples with N-stage 1 were stratified (log rank, p= 3e-05) in high/low risk groups with HR= 21.11 and p-value= 0.0046. (f) Patients in T stage 4 were stratified (log rank, p= 0.001) in high/low risk groups with HR= 13.65 and p-value= 0.0124.

### Web-server for risk prediction in CRC patients: CRCRpred

We also developed a web-server CRCRpred, which is freely available at https://webs.iiitd.edu.in/raghava/crcrpred, in order to provide service to the community. This web-server implements the current study in order to predict high and low risk patients given the nanomolar concentration of required Bcl2 family protein(s). Following are the two prediction modules with their brief descriptions:

#### (i) Single-protein prediction

Often the user will not possess the concentrations of all required Bcl2 family proteins, mainly because protein level quantification is a challenging task in itself. Keeping this in mind we provide this module to the user where, with the limited knowledge of concentration(s) of one or more than one proteins, the risk can be predicted. The prediction here is a protein wise prediction. Input concentration from the user is fed into a linear regression model and risk percentage is estimated. This regression model comprises of fitting bin-wise mean protein concentrations with probability of high risk patients in that bin. High risk and low risk patient stratification was done on the basis of median survival time in CRC dataset.

#### (ii) Multiple-proteins prediction

This module computes the Risk Score (RS) of a patient, given the concentrations of all five proteins by estimation of RS for the given patient. The patient is classified into high/low risk category based on the cutoff, RS=0. The distance from cutoff point is displayed as percentage risk to the user along with the risk grade.

## Discussion

Colorectal cancer is a mortal disease, with incidences all over the world, and requires better clinical strategies. Recent experiments have suggested a crucial role of apoptotic type 2 pathway in tumor progression and development, as a result of which many new drugs are targeted on the proteins involved in this pathway. For e.g. several BH3 mimetics, which are small molecules that target and inhibit anti-apoptotic Bcl2 proteins, such as ABT-737, ABT-263 (Navitoclax), ABT-199 (Venetoclax), WEHI-539, A-1155463 etc. have been introduced in the market while several others are in clinical trials [22–24]. Thus, monitoring the protein profile in this pathway is deemed to be a good strategy in order to identify high and low risk patients in a post-diagnosis-pre-therapeutic setting, which could then help to estimate the risk at hand and decide between the existing remedies to go with. However, the trend of this protein concentration profile in the case of CRC patients does not correlate often, partly due to variation in expression of functional paralogs and/or genetic/epigenetic changes. This alteration in the protein profile is one of the major reasons for chemotherapeutic failures (relapses and deaths). Hence, restricting to a specific marker protein (for e.g BclXL alone) to identify high/low risk CRC cases might not be a good way to solve this problem. We make this evident here by performing a two variable t-test between protein concentrations in favorable and unfavorable patient cases, which showed significant difference in Bak, Bax, BclXL and Mcl1 levels. However, it is for certain that the death pathway is defunct and there exists an unexploited relationship between the overall protein profile and survival of the cancer patient.

To tackle this, we first took at the total pro- and anti-apoptotic protein concentration, since both of these are upregulated in the event of cellular stress conditions such as tumor [10], and stratified the patients on the basis of mean and median cutoffs of this total sum. We also took the difference in the levels of these and repeated the same procedure. While, the former was able to differentiate high and low risk CRC patients effectively (HR value higher than DR MOMP), the latter resulted in an insignificant p-value at median cutoff. Next, we constructed multi variable linear regression models using five-fold cross validation and implementing various techniques. The predicted OS from one of these techniques (LassoLars) was found to stratify high and low risk patients with a high HR, but still performed poorly in comparison to BclXL alone. We then analysed linear combinations of Bcl2 family proteins by making use of a five-fold parameter optimization technique and constructed a parameter Risk Score (RS) which is a remnant of altered protein profile (including paralogs) and/or binding affinities. We found that RS outperforms the task of risk stratification as compared to previous studies as evident on the basis of risk estimates and KM plots. RS elucidates the significance of anti-apoptotic protein concentration in cancer patients as observed from the negative *β\** value obtained for each patient. Patients with RS*<*0 (mean cut-off) or RS*<*0.266 (median cut-off) are classified as high risk patients, indicating that a higher anti-apoptotic concentration increases the risk associated with CRC. This has a direct correlation with cell death mechanism related to type 2 apoptotic pathway, where a higher anti-apoptotic concentration as compared to pro-apoptotic concentration has been shown to be the reason for evasion of apoptosis by the tumor cell [25]. RS also elucidate the importance of BclXL and Mcl1 as biomarkers for risk assessment, as is evident by the coefficients, over other proteins which has also been noted in the previous studies [16, 17, 26–29]. We demonstrate the power of RS by prediction of favorable/unfavorable outcomes, with known protein concentrations of Bcl2 family proteins. A comparison was shown with a recent prognostic marker (DR MOMP), to establish the superiority of RS. We also looked at the significance of RS when other pathological features are taken into account, and noticed that in a multivariate analysis, it could stratify patients with upto 29 fold mortality risk than low risk ones. RS was also able to stratify high and low risk CRC patients in sub-classes formed on the basis of pathological factors which is shown here by KM plots and logrank tests thereby emphasizing the significant differences between the two classified groups for each of these features. It should be noted that these sub-groups are claimed to be prone to high risk of death by previous studies and surveys [18–21]. A prognostic marker which could classify patients among these sub-groups could be very beneficial for deciding the fate of the patients and future therapy. RS could be one such candidate marker.

To provide valuable help to scientific community, a web server, CRCRpred, https://webs.iiitd.edu.in/raghava/crcrpred was also developed. CRCRpred is built on a responsive template, compatible for desktop, tablet, and smartphone. These templates are dynamic that fit content based on screen size of the device. This web server can be used to distinguish high risk CRC patients from low risk CRC patients given the protein concentration of one or more apoptotic proteins (Bak, Bax, Bcl2, BclXL or Mcl1) involved in the process of Mitochondrial Outer Membrane Permeabilization (MOMP), which is defunct in the case of cancer. This risk estimation is based on statistical and survival analysis on a recent CRC dataset and risk score (RS) obtained from the same can be a promising prognostic tool in clinical/research domain.

## Supporting information

**S1 File Additonal results.** KM plots and multivariate analysis results.

**S2 Table 1 Clinical dataset with predicted survival times** This table contains the clinical information about the patients used in this study as adapted from Lindner AU, et al. Gut. 2017. OS predicted from multiple regression models are also provided.

**S2 Table 2-7 Estimated** *β*^*i*^ **for various optimized parameters** *W* ^*i*^ The tables contain estimated *β* values for specified weight parameter W, for each patient. Risk Score (RS) values (z-normalized *β*^*\**^) are provided in Table 7.

## Acknowledgments

Authors are thankful to funding agencies Department of Science and Technology (J. C. Bose Fellowship), India; University Grants Commission (UGC), Govt. of India and Indraprastha Institute of Information Technology for financial support and fellowships.

